# MS-driven metabolic alterations are recapitulated in iPSC-derived astrocytes

**DOI:** 10.1101/2021.08.27.457853

**Authors:** Bruno Ghirotto, Danyllo F. Oliveira, Marcella Cipelli, Paulo J. Basso, Jean de Lima, Cristiane N. S. Breda, Henrique C. Ribeiro, Camille C. C. Silva, Meire I. Hiyane, Elia G. Caldini, Alessandra Sussulini, Alicia J. Kowaltowski, Enedina M. L. Oliveira, Mayana Zatz, Niels O. S. Câmara

## Abstract

**Objective:** Astrocytes play a significant role in the pathology of Multiple Sclerosis (MS). Nevertheless, for ethical reasons, most of the studies in these cells were performed on the Experimental Autoimmune Encephalomyelitis model. As there are significant differences between human and mouse cells, we aimed here to better characterize astrocytes from patients with MS (PwMS), focusing mainly on mitochondrial function and cell metabolism.

**Methods:** We obtained and characterized induced pluripotent stem cell (iPSC)-derived astrocytes from three PwMS and three unaffected controls and performed functional assays including electron microscopy, flow cytometry, cytokine measurement, gene expression, *in situ* respiration, and metabolomics.

**Results:** We detected several differences in MS astrocytes including: (i) enrichment of genes associated with mitophagy and neurodegeneration, (ii) increased mitochondrial fission and decreased mitochondrial to nuclear DNA ratio, indicating disruption of mitochondrial content, (iii) increased production of superoxide and MS-related proinflammatory chemokines, (iv) increased electron transport capacity and proton leak, in line with the increased oxidative stress, and (v) a distinct metabolic profile, with a deficiency in amino acid catabolism and increased sphingolipid metabolism, which have already been linked to MS.

**Interpretation:** To our knowledge, this is the first study thoroughly describing the metabolic profile of iPSC-derived astrocytes from PwMS, and validating this model as a powerful tool to study disease mechanisms and to perform non-invasive drug targeting assays *in vitro*. Our findings recapitulate several disease features described in patients and provide new mechanistic insights into the metabolic rewiring of astrocytes in MS, which could be targeted in future therapeutic studies.

## INTRODUCTION

Multiple sclerosis (MS) is a multifactorial demyelinating disease of the central nervous system (CNS) that affects more than two million people worldwide**^1^**, characterized by neurological deficits that usually arise when the patients are around 30 years old. It is clinically heterogeneous and can include sensory, vision, and motor problems, as well as fatigue, pain, and cognitive impairments**^2^**. Although many therapies that alleviate MS symptoms have been developed, there is still no cure**^3^**. Patients usually start presenting symptoms that are consistent with relapsing-remitting MS (RRMS), in which neuronal deficiencies are recurrent but reversible, and may transition later to secondary progressive MS (SPMS), when neurological damage becomes progressive and irreversible**^3^**.

Several studies have indicated that astrocytes, the major cell population in the human brain, play a key role in the pathology of MS, enhancing neuroinflammation and oxidative stress in the CNS**^4^**. Interestingly, a recent study**^5^** identified mitochondrial dysfunction as a pathway in astrocyte pathology both in MS and its animal model of Experimental Autoimmune Encephalomyelitis (EAE). Accordingly, mitochondrial dysfunction has already been linked to several neurodegenerative diseases**^6,7^**.

In contrast to the abundant literature available in EAE, the number of studies analyzing the reactivity of astrocytes in patients with MS (PwMS) are limited, mainly due to the impossibilities to obtain CNS biopsies. There are significant known differences between human and mouse astrocytes at the baseline level and upon inflammatory stimuli**^8^**. Another relevant aspect to be considered is that the heterogeneity observed in MS is poorly represented in animal models**^9^**. In this context, the use of human induced pluripotent stem cells (iPSCs) emerges as a powerful tool to study MS molecular mechanisms in CNS-resident cells. The generation of iPSCs from affected patients with different clinical presentations allows for the discovery of molecular mechanisms that may be responsible for the transition to the most severe forms of the disease, thus uncovering new possible therapeutic avenues**^9^**.

Here we obtained iPSC-derived astrocytes from three RRMS patients and three unaffected controls and thoroughly investigated mitochondrial function and metabolism, parameters that had been described superficially in this human cellular model. We found several changes in patient cells regarding mitochondrial morphology, respiration, and cell metabolism, along with chemokine and superoxide production. Our findings bring substantial novel insights to the characterization of patient astrocytes in the context of MS, and open new perspectives in terms of disease modeling and drug targeting studies in the field.

## METHODOLOGY

### Ethical compliance

All human procedures in this study and the consent forms were approved by the Ethics Committees on Human Research of the Institute of Biomedical Sciences – University of São Paulo (CAAE n° 26052519.9.0000.5467), of the Institute of Biosciences – University of São Paulo (CAAE n° 26052519.9.3001.5464), and of the Federal University of São Paulo (CAAE n° 26052519.9.3002.5505).

### Generation and maintenance of human iPSCs

Human iPSCs were generated from peripheral blood mononuclear cells (PBMC) obtained from three RRMS patients and three age and sex-matched controls after signed consent. The clinical data of the patients are detailed in Table 1. PBMC reprogramming and iPSC culture maintenance were performed as previously described**^10^**. One iPSC clone was selected from each subject for further characterization and analysis.

**Table 1.** Clinical data of the subjects included in this study.

| Subjects | Age (y) | Gender | Diagnosis | Relapses | First Symptom | OCBs | Medication | Time of Symptoms |
| --- | --- | --- | --- | --- | --- | --- | --- | --- |
| MS1 | 61 | M | RRMS | 10 | Brainstem | Positive | None | 29 years |
| MS2 | 35 | F | RRMS | 1 | Spine | N/A | IFNβ-1a | 4 years |
| MS3 | 56 | F | RRMS | 4 | Sensitive | N/A | None | 21 years |
| CTL1 | 63 | M | Control | N/A | N/A | N/A | N/A | N/A |
| CTL2 | 52 | F | Control | N/A | N/A | N/A | N/A | N/A |
| CTL3 | 57 | M | Control | N/A | N/A | N/A | N/A | N/A |

**Legend:** M: male; F: Female; OCB: oligoclonal bands of immunoglobulin G in the cerebrospinal fluid; IFN $\beta$ -1a: interferon beta 1a; y: years; N/A: not available; MS: Multiple Sclerosis; CTL: unaffected controls.

### Multiplex ligation-dependent probe amplification (MLPA) assay

The total genomic DNA was extracted from iPSCs using the DNeasy Blood and Tissue kit (QIAGEN, Germany), according to the manufacturer’s instructions. MLPA analysis was performed using subtelomeric kits (P036 and P070; MRC - Holland). The PCR products were detected by the ABI3130 Genetic Analyzer (Applied Biosystems, USA) using capillary electrophoresis and the raw data generated was analyzed with GeneMarker software (Softgenetics, USA).

### Differentiation of iPSCs into neural progenitor cells (NPCs) and astrocytes

NPC and astrocyte differentiations were performed as described**^10^**. Cell viability was analyzed with Trypan-Blue (Gibco, USA) staining using the Countess automated cell counter (Thermo, USA). The cells were tested for mycoplasma contamination using the MycoAlert kit (Lonza, Switzerland). An overview of our differentiation procedure is shown below (Fig 1).

**Figure 1.**
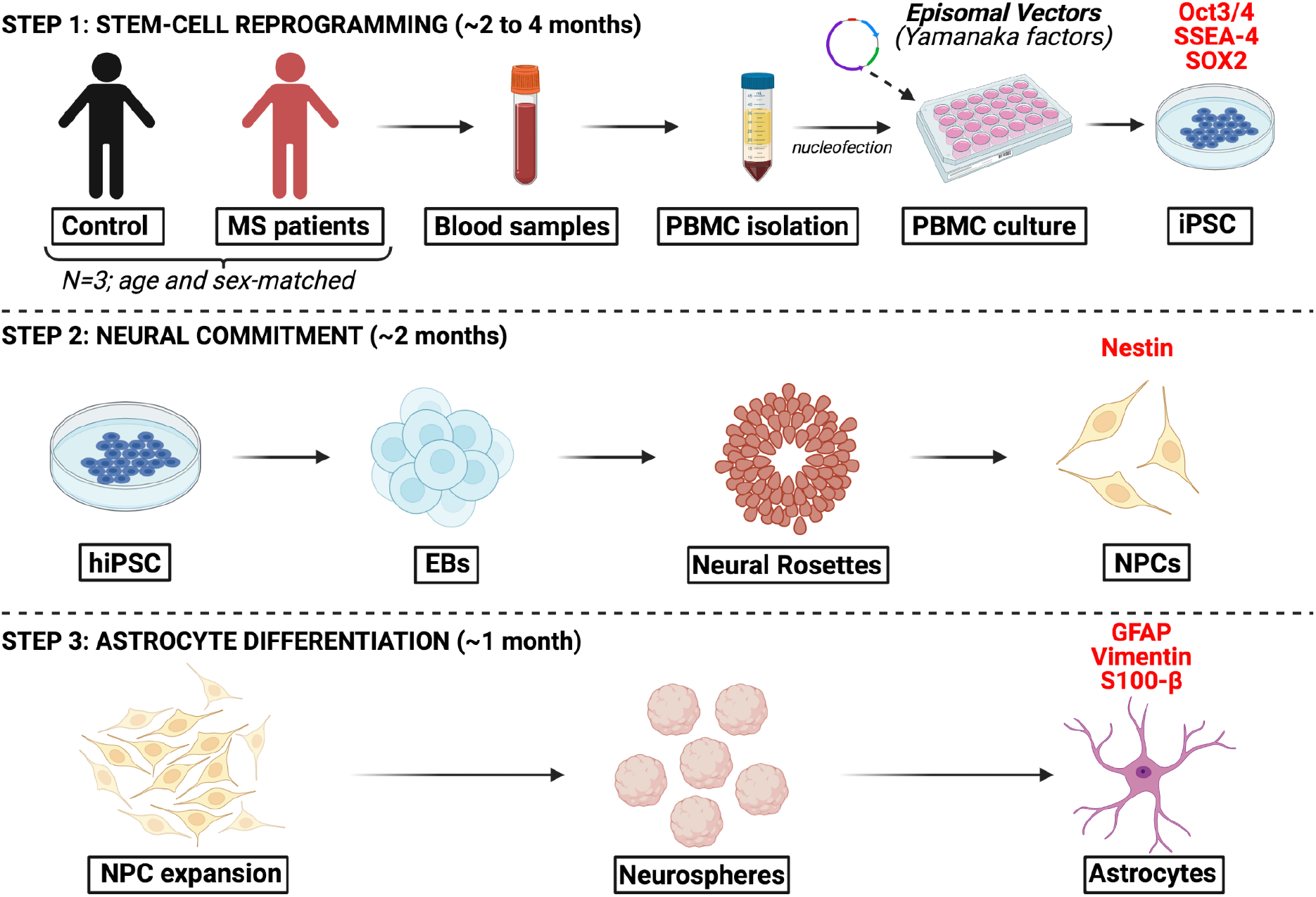
Experimental Design. The cell culture protocol consists of 3 main steps: **Step 1**, Stem-cell reprogramming, requires 2 to 4 months to be completed. Blood samples were obtained from 3 PwMS and 3 age- and sex-matched controls, and used for isolation of PBMCs. The PBMCs were cultured and nucleofected with episomal vectors containing the Yamanaka factors required for cell reprogramming into iPSCs. To characterize the iPSCs, we used Oct3/4, SSEA-4 and SOX2 markers; **Step 2**, Neural commitment, lasts approximately 2 months. iPSCs were cultured in suspension as EBs, exposed to neural differentiation factors and then plated to form neural rosettes. Rosettes were manually picked, expanded and dissociated, giving rise to the NPCs. Here, we characterized NPCs as Nestin^+^ cells; **Step 3**, Astrocyte differentiation, takes about 1 month and consists of NPC expansion and culture in suspension as neurospheres, exposed to astrocyte differentiation factors. Neurospheres are then plated and give rise to astrocytes, which were the cells used for further experiments in this study. We used GFAP, Vimentin and S100-β as astrocyte markers. The figure was generated using Biorender.

### PCR Array

Total RNA was isolated from astrocytes using the RNeasy mini kit (QIAGEN, Germany), together with on-column DNAse digestion using the RNAse-free DNAse set (QIAGEN, Germany). cDNA was synthesized from 500ng RNA, using the RT2 First Strand Kit (QIAGEN, Germany). PCR array reaction was performed using the Human Mitochondria PCR array (QIAGEN, PAHS-087Z, Germany), following the manufacturer’s instructions. The reactions were performed on the QuantStudio 12K Flex Real-Time PCR System device (Applied Biosystems, USA). Data analysis was performed on the GeneGlobe software (QIAGEN, Germany), being the beta-2-microglobulin gene (*B2M*) chosen to normalize all the samples. The CT threshold was standardized throughout the plates, as recommended by the manufacturer. The fold regulation threshold used during the analysis was 1.5. Genes with undetermined CTs in at least one sample were excluded from the analysis. Enrichment analysis of the upregulated genes was performed with the EnrichR software**^11^** using the KEGG pathways and GO Biological Process databases.

### Transmission electron microscopy

To perform electron microscopy assays, 1×10^6^ astrocytes were treated with Accutase (Gibco, USA), centrifuged, and fixed in a 2% glutaraldehyde solution (Ladd Research Industries, USA). The cell pellet was embedded in a specific resin and carefully placed on copper membranes for subsequent analysis by transmission electron microscopy using a Jeol 1010 (Tokyo, Japan) electron microscope. At least 10 individual cells per group were analyzed. Mitochondrial morphology was assessed using the ImageJ software, as described**^12^**, through the quantification of automated mitochondrial aspect ratio and mitochondrial roundness. Additionally, cell areas were individualized and for each image the ratios between the area of individual mitochondria/area occupied by cell were determined.

### Oxygen consumption and extracellular acidification rates (OCR and ECAR)

We first seeded 40000 astrocytes per well on Matrigel (Corning, USA)-coated Seahorse XF24 cell plates (Agilent, USA) 2 days before the experiment in ABM medium plus supplements (Lonza, Switzerland). Before measurements, cells were washed 3 times and incubated for 1 h at 37 °C in an incubator without CO_2_, with respiratory medium (pH 7.4) consisting of DMEM (Gibco, USA) supplemented with 25 mM glucose (Sigma, USA) plus 1 mM pyruvate (Sigma, USA) and 2 mM glutamine (Gibco, USA). After equipment calibration, baseline respiration measurements were followed by injection of 1 μM oligomycin (Sigma, USA), 500 μM 2,4-dinitrophenol (2,4-DNP) and 1 μM rotenone plus 1 μM antimycin A. Concentrations of modulators were determined through preliminary titration experiments (not shown). To perform the glycolysis stress test, we used Seahorse XF minimal DMEM medium (Agilent, USA) supplemented with 1 mM glutamine (Gibco, USA). The injections consisted of 10mM glucose (Sigma, USA), 1 μM oligomycin (Sigma, USA) and 50 mM 2-deoxi-D-glucose (2-DG) (Sigma, USA). All respiratory modulators were previously titrated, and all plates were normalized to protein content using the BCA kit (Thermo, USA). Analysis was performed using the Wave software (Agilent, USA).

### Immunofluorescence

iPSC, NPC and astrocyte cultures were first plated in Matrigel (Corning, USA)-coated Lab-tek 8-well chamber slide systems (Thermo Fisher, USA) at a density of 4×10^4^ cells per well, then fixed with 4% PFA, followed by permeabilization with 0.01% Triton X-100, and blocking with 5% BSA in PBS. The cells were then incubated overnight with the following primary or conjugated antibodies: rabbit anti-OCT4 (ab19857, Abcam, 1:100); APC anti-SSEA4 (FAB1435A, R&D systems, 1:100); AF647 anti-nestin (560393, BD, 1:100); mouse anti-GFAP (G3893, Sigma, 1:100); mouse anti-Vimentin (M7020, DAKO, 1:100) at 4 °C, and subsequently incubated with the following secondary antibodies: AF488 goat anti-rabbit IgG (A11034, Life, 1:1000); AF488 goat anti-mouse IgG (A11001, Life, 1:250); AF594 donkey anti-mouse IgG (a21203, Life, 1:250) for 1 h at room temperature. Slides were mounted with Vectashield containing DAPI (Vector Laboratories, USA), dried, sealed, and the analysis was performed using a Zeiss LSM 800 confocal microscope. Quantification of the percentage of positive cells was performed using ImageJ software.

### Mitochondrial superoxide production

To estimate the mitochondrial superoxide production, we used flow cytometry. Astrocytes were initially treated with Accutase (Gibco, USA), washed with 1x PBS, centrifuged at 1500 rpm at room temperature for 5 minutes and then resuspended in a staining solution of PBS, 5 μM of MitoSOX and 0.2% LIVE/DEAD, all from Life, USA. Then, cells were incubated at 37 °C for 20 minutes, centrifuged and resuspended in PBS for analysis. Flow cytometry was performed on a BD FACSCanto™ II Cell Analyzer (BD, USA). For each sample, we acquired 200000 events. Analysis of frequency and mean fluorescence intensity (MFI) of the probes in the samples was performed using the FlowJo software (Flow Jo, USA). Gating was performed in single cells/live cells/positive populations for each marker.

### Cytokine production

Proinflammatory cytokines CCL5, G-CSF, CXCL-10, GM-CSF, and IL-6 were quantified in astrocyte culture supernatants using the Bio-Plex Pro Human Cytokine kit (Biorad, USA), according to the manufacturer’s instructions. Astrocytes were plated at a density of 5×10^5^ cells/well and activated with 20ng/mL recombinant human TNF-α (Peprotech, USA) for 48h prior to cytokine measurements.

### Western-blotting

Astrocytes were lysed by RIPA buffer and approximately 50 μg of total protein was diluted in sample buffer (Laemmli-BioRad, USA) containing 20 mg/mL of DTT (Sigma-Aldrich, USA). Proteins were denatured by heating (5 min at 95 °C), and then separated by electrophoresis in a 10% polyacrylamide gel. Subsequently, the proteins were transferred to a nitrocellulose membrane, and blocked for 1 hour in 5% milk dissolved in Tris-Buffered Saline (TBS) containing 0.05% of Tween-20 (TBS-T) for further incubation with primary antibodies overnight. After this incubation, the membrane was washed with TBS-T and incubated for 1 hour with secondary antibodies conjugated with Horseradish peroxidase (HRP). The molecular mass of the proteins was determined by comparison with the migration pattern of Spectra Multicolor Broad Range Protein Ladder (Thermo, USA). Membranes were developed with SuperSignal Chemiluminescent Substrate (Thermo, USA) and read using the LAS-500 equipment (GE, USA). Primary antibodies used were rabbit anti-DRP1 (ab184247, Abcam, 1:1000) and mouse anti-β-actin (A5441, Sigma, 1:10000). Secondary antibodies used were anti-mouse (A9044, Sigma, 1:5000) and anti-rabbit (SAB3700861, Sigma, 1:5000).

### RNA extraction, cDNA synthesis and RT-qPCR

Total RNA was isolated from iPSCs, NPCs, and astrocytes using the RNeasy mini kit (QIAGEN, Germany). cDNA was synthesized from 2000ng RNA using the M-MLV Reverse Transcriptase System (Promega, USA). The qPCR reaction was performed with 0.5 μL (500 nM) of the forward primer, 0.5 μL (500 nM) of the reverse primer, 1 μL of purified water, 5 μL of SYBR Green Master Mix (Applied Biosystems, USA) and 3 μL of the previously diluted cDNA. All reactions were performed on the QuantStudio 12K Flex Real-Time PCR System device (Applied Biosystems, USA). All samples were analyzed in technical triplicates, and the *GAPDH* or *RPLP0* genes were chosen as housekeeping. Quantification was performed by the 2-ΔΔCT method. We used the following primer sequences: *OCT4* (F: TCCCATGCATTCAAACTGAGG; R: CCAAAAACCCTGGCACAAACT); *NANOG* (F: TGGACACTGGCTGAATCCTTC; R: CCAAAAACCCTGGCACAAACT); *SOX2* (F: GCTACAGCATGATGCAGGACCA; R: TCTGCGAGCTGGTCATGGAGTT); *GAPDH* (F: ACAACTTTGGTATCGTGGAAGG; R: GCCATCACGCCACAGTTTC); *S100B* (F: GAAGAAATCCGAACTGAAGGAGC; R: TCCTGGAAGTCACATTCGCCGT); *16S* (F: GCCTTCCCCCGTAAATGATA; R: TTATGCGATTACCGGGCTCT) and *RPLP0* (F: CCTCATATCCGGGGGAATGTG; R: GCAGCAGCTGGCACCTTATTG).

### UHPLC-MS metabolomics

Metabolite extraction consisted of the addition of an 80% methanol solution to 2×10^6^ astrocytes, followed by overnight drying of the pellets in a vacuum concentrator (Eppendorf, Germany). For reverse phase chromatography, the samples were analyzed on an Ultimate 3000 (Thermo, Germany) system equipped with an Acquity CSH C18 (2.1 × 100 mm, 1.7 μm) column (Waters, USA) at 35°C and a 27 min chromatographic run was performed with a 0.3 mL/min flow and an injection volume of 2 μL. The mobile phases consisted of 0.1% (v/v) formic acid in water and 0.1% (v/v) formic acid in acetonitrile (B). The following gradient was used: 5% B for 15 min, 98% B for 7 min, and 5% B for 5 min. For hydrophilic interaction chromatography (HILIC), the samples were analyzed by the same instrument with a Acquity BEH Amide (2.1 × 100 mm, 1.7 μm) column (Waters, USA) at 35°C and a 15 min chromatographic run was performed with a 0.3 mL/min flow and injection volume of 3 μL. The mobile phases consisted of 0.1% (v/v) formic acid in 10 mM ammonium formate (A) and 0.1% (v/v) formic acid in (90:10) acetonitrile: 10 mM ammonium formate (B). The following gradient was used: 100% B for 9 min, 50% B for 3 min, and 100% B for 3 min. Mass spectrometry analyses were performed on a Q-Exactive mass spectrometer (Thermo, Germany), operating in positive and negative ionization modes. The spray voltage was 3.5 kV, the capillary temperature was 256°C, the auxiliary gas temperature was set to 413 °C with a flow rate of 11 L/min, and the sweep gas and desolvation gas flow rates were 2 and 48 L/min, respectively. RF lens were set at 50 V. A mass range of *m/z* 100-1500 was used for full scan mode. MS/MS precursor selection was set for the most intense 5 ions (DDA mode) with an isolation window of *m/z* 4.0. We performed batch analysis with quality control (QC) samples, which consisted of a pool of all the samples included in the study to avoid possible sources of variation. The QC samples were injected five times at the beginning of the batch, after every set of 3 injections and at the end of the batch. The raw MS data was transformed using abf File Converter (www.reifycs.com/AbfConverter) for data preprocessing. Data preprocessing, statistical analysis, and visualization were performed using MetaboAnalyst**^13^** and R programming language**^14^**. Peak detection, deconvolution, alignment, identification, compound annotation, and background subtraction were performed using the MS-Dial software. Compound annotation was performed using LipidBlast**^15^** and MassBank of North America library for polar metabolites, and manually curated to prevent false positives/negatives. Only metabolites with a similarity score higher than 70% were selected. After preprocessing, the data were normalized using LOWESS normalization on MS-Dial and the normalized data was exported for statistical analysis. A relative standard deviation cutoff of 30% was applied on the dataset, followed by log2 transformation and autoscaling. Quality assessment was performed by principal component analysis (PCA) on MetaboAnalyst**^13^**. *‘Barplot’*, *‘heatmap.2’*, and *‘gplot’* libraries and packages were used for data visualization. To observe the differential metabolites between the samples, variable selection methods were applied. Each dataset was subjected to *t*-test and PLS-DA VIP score analysis using the *‘stats’* and *‘mixOmics’* packages. Metabolites with VIP score > 1.0 and a *p*-value < 0.05 were selected for further analysis.

### Statistical analyses

Differences between groups were first analyzed using the Kolmogorov-Smirnov normalization test. If both groups tested positive, unpaired two-tailed Student’s *t*-test was used to compare the groups. Otherwise, the nonparametric Mann-Whitney test was selected. The *p*-values < 0.05 were considered significant. All graphs were generated using the GraphPad Prism Software.

## RESULTS

### Characterization of the subjects

We obtained samples from three patients with RRMS and three age and sex-matched controls. The clinical data of these subjects included in our study is summarized below (Table 1).

### Characterization of iPSCs, NPCs and astrocytes

We reprogrammed iPSCs from PBMCs of control and PwMS successfully. All cells were positive for the pluripotent markers OCT4 (nuclear) and SSEA-4 (surface) (Fig 2A) and did not present copy number variations, neither duplications or deletions, in chromosomes, (Fig 2B), in the MLPA analysis. We then analyzed the expression of the pluripotency genes *SOX2* (Fig 2C), *OCT4* (Fig 2D) and *NANOG* (Fig 2E) and observed no significant differences between the control and MS groups. The NPCs were all positive for the Nestin marker (Fig 2F), with no differences in the frequency of positive cells between control and MS groups (Fig 2G). Astrocytes were positive for GFAP and Vimentin in both groups (Figure 2H), with no differences in the frequency of GFAP and Vimentin (Fig 2I-J)-positive cells between them. Additionally, no differences in the expression of the *S100B* gene were observed between the groups (Fig 2K).

**Figure 2.**
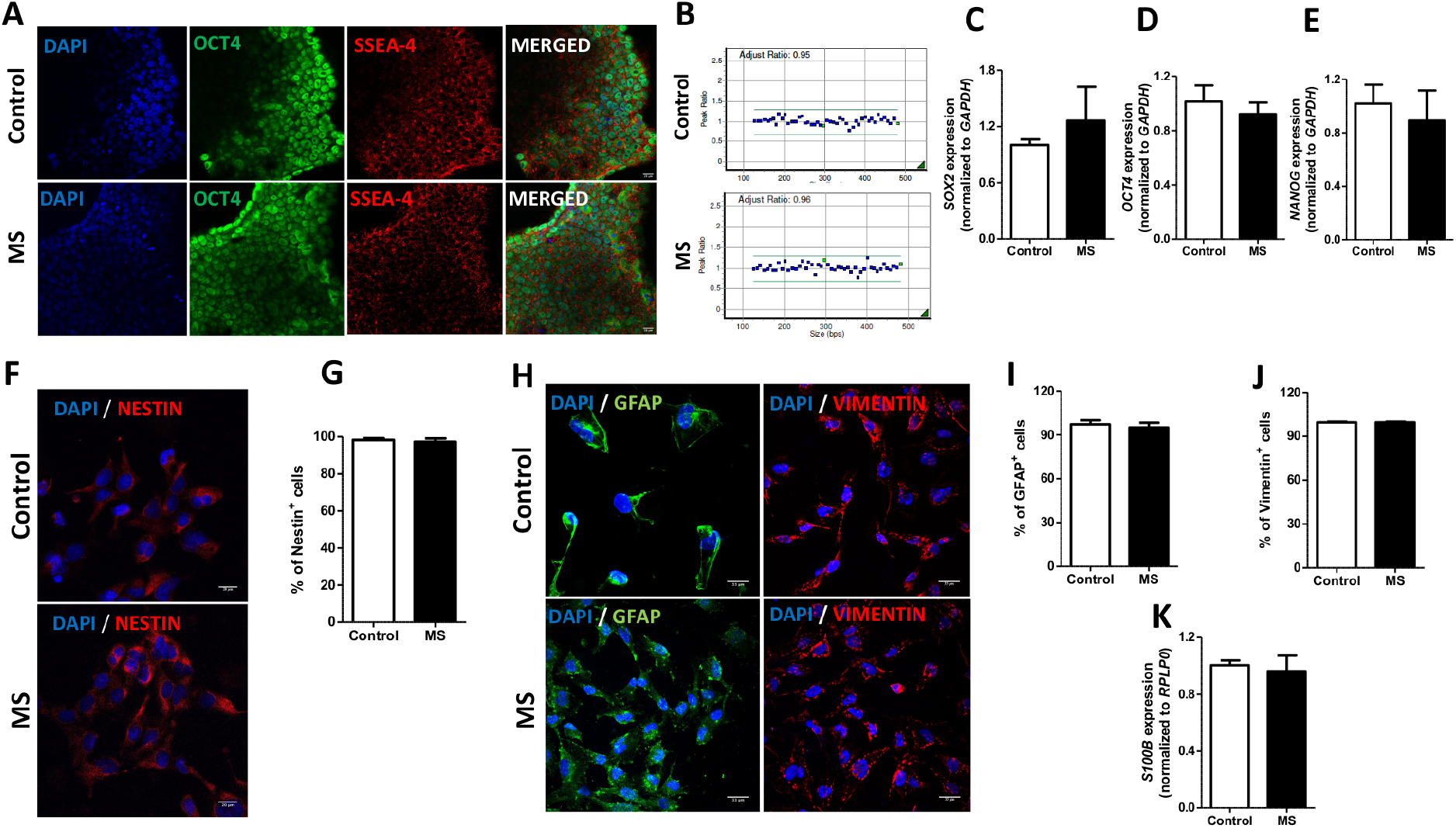
iPSC, NPC and astrocyte characterization. **A.** Panel of confocal microscopy analysis for the pluripotency markers OCT4 (green), SSEA-4 (red) and DAPI (blue), for nuclear staining. The top line is representative of the results obtained for the control group and the bottom line is representative of the results obtained for the MS group. Scale bar = 20μm. **B.** Representative MLPA analysis for the control and MS groups using the P036 and P070 kits (MRC – Holland). **C-E.** Expression analysis of the pluripotency genes *SOX2* **(C)**, *OCT4* **(D)** and *NANOG* **(E)** using RT-qPCR, normalized to *GAPDH*. **F.** Confocal microscopy analysis for the NPC marker Nestin (red) and DAPI (blue) for nuclear staining. The top panel is representative of the control group and the bottom panel is representative of the MS group. Scale bar = 20μm. **G.** Quantification of the frequency (%) of Nestin positive cells between the groups using the ImageJ software. **H.** Confocal microscopy analysis for GFAP (green), Vimentin (red) and DAPI (blue) for nuclear staining. The top line is representative of the results obtained for the control group and the bottom line is representative of the results obtained for the MS groups. Scale bar = 33μm. **I-J** Quantification of the frequency (%) of GFAP **(I)** and Vimentin **(J)** positive cells between the groups using the ImageJ software. **K.** Expression analysis of the *S100B* astrocyte marker using RT-qPCR, normalized to *RPLP0*. N=3 controls and 3 patients, analyzed in technical triplicates in two independent experiments. Data are represented as mean ± standard error.

### MS astrocytes show altered expression of mitochondrially-related genes

Metabolic and mitochondrial functional alterations are intimately linked with a variety of neurological and inflammatory diseases, including MS**^6, 7^**. To gain insight into mitochondrial biology in MS versus control astrocytes, we performed a PCR array analysis of human mitochondria-related genes. An upregulation of 18 genes was uncovered in our analysis, as well as a downregulation of one gene in astrocytes derived from PwMS compared to controls (Fig 3A). We performed an enrichment analysis on the 18 upregulated genes shown in Figure 3A using the KEGG database and observed a significant enrichment of 8 pathways, related mostly to cell death, mitochondrial dysfunction and neurodegeneration (Fig 3B). To have further insights on the processes regulated, we performed a biological processes enrichment analysis on the same gene list using the GO database and observed a significant enrichment of terms related to mitochondrial transport and cell death (Fig 3C). These results indicate that MS astrocytes have a different expression of mitochondrially-related genes and therefore should display altered mitochondrial morphology and function.

**Figure 3.**
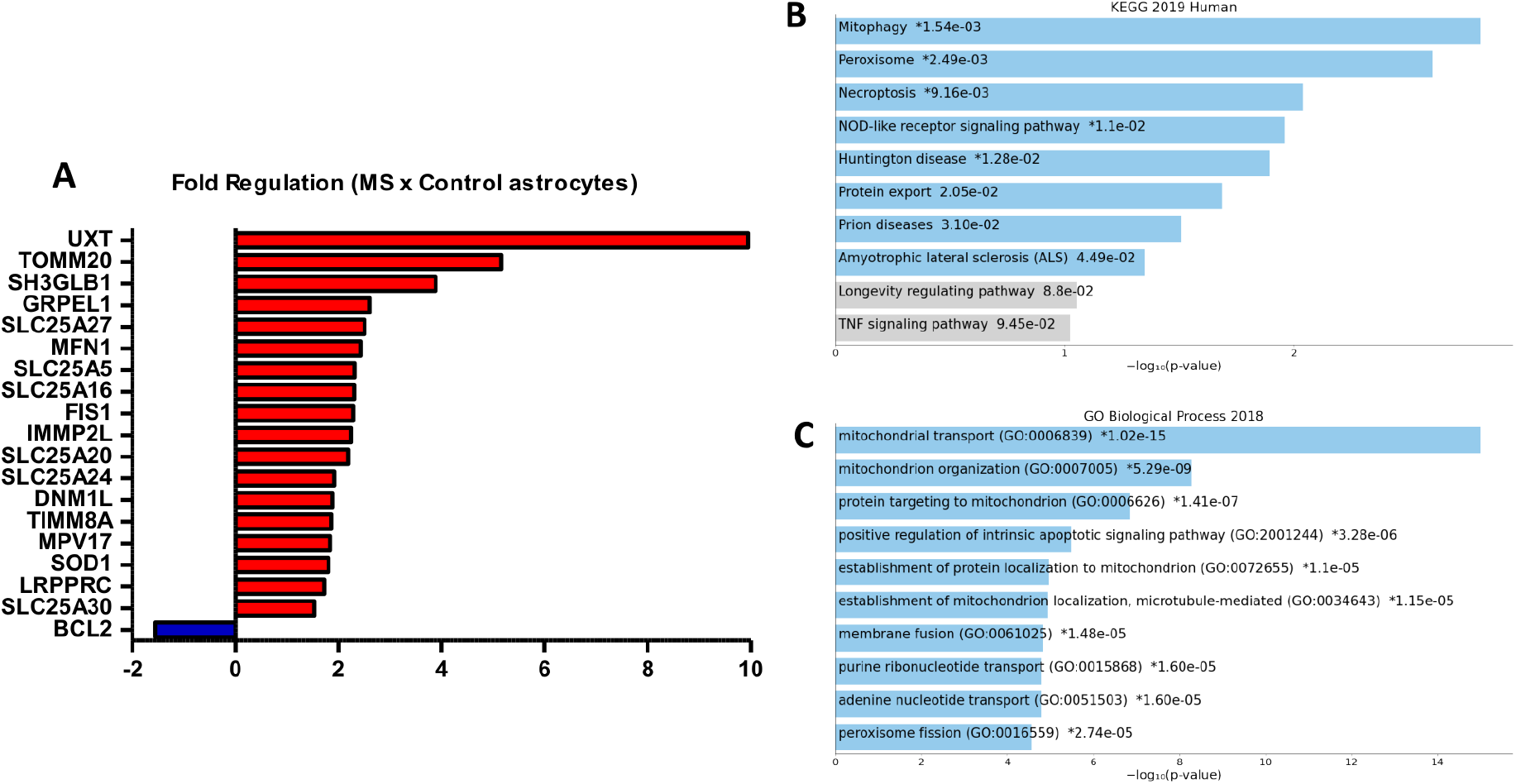
PCR array analysis. **A.** Bar graph of absolute fold regulation values of genes in MS astrocytes, compared to the control group. Red bars and blue bars represent genes above and below the fold regulation threshold (1.5), respectively. N=3 controls and 3 PwMS. Expression was normalized to the *B2M* housekeeping gene. Fold regulation values represent fold-change results in a biologically meaningful way. **B-C** Enrichment analysis was performed on the 18 top genes shown in panel A using the KEGG 2019 human database **(B)** and the GO Biological Process 2018 database **(C)**. The bar charts show the top 10 enriched terms in the chosen library, along with their corresponding *p*-values. Colored bars correspond to terms with significant *p*-values (<0.05). An asterisk (*) next to a *p*-value indicates the term also has a significant adjusted *p*-value (<0.05). Analyses were performed using the EnrichR software.

### MS astrocytes have increased mitochondrial fission

Increased mitochondrial fission in resident CNS cells, such as astrocytes and microglia, propagates neuroinflammation and neurodegeneration**^16^**. To evaluate mitochondrial morphology, we used transmission electron microscopy (Fig 4A) and observed that MS astrocytes display increased mitochondrial roundness (Fig 4B) together with decreased aspect ratio (Fig 4C, a measure of elongation), which is consistent with mitochondrial fragmentation**^12^**. In this sense, we also observed increased protein levels of DRP1, which is a key regulator of the mitochondrial fission machinery, in MS astrocytes (Fig 4D-E). Additionally, we observed a decreased ratio between the mitochondrial area and the area occupied by cells in MS astrocytes (Fig 4F), which is indicative of decreased mitochondrial mass. We then assessed the mitochondrial/nuclear DNA ratio using RT-qPCR and observed a significantly decreased ratio in the MS astrocytes (Fig 4G), confirming decreased mitochondrial content in these cells.

**Figure 4.**
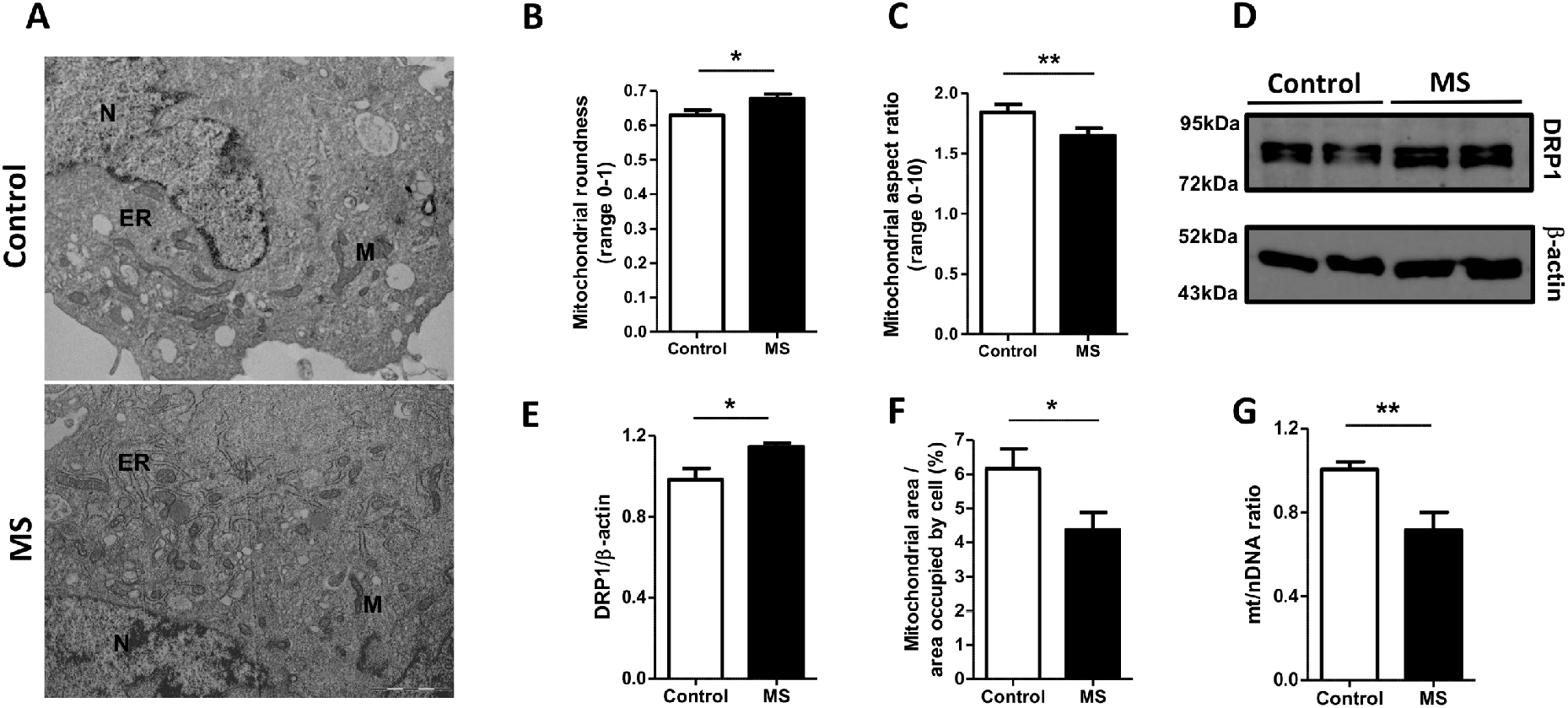
Increased mitochondrial fission in MS astrocytes. **A.** Electron microscopy analysis. The top line is representative of the results obtained for the control group cells and the bottom line is representative of the results obtained for the MS group cells. Scale bar = 2μm. N: nucleus, ER: endoplasmic reticulum, M: mitochondria. **B.** Quantification of mitochondrial roundness in control and MS astrocytes. **C.** Quantification of mitochondrial aspect ratio in control and MS astrocytes. For panels **A-C**, N=10 individual cells per group and quantification was performed using the ImageJ software. **D-E.** Representative Western-Blot membrane **(D)** and quantification **(E)** for DRP1 (top panel, predicted molecular weight = 83kDa) and β-actin (bottom panel, predicted molecular weight = 45kDa). The first two samples in the membrane are from the control astrocytes and the last two samples are from the MS cells. These data are representative of cells from N=2 controls and 2 patients from two independent experiments. Quantification was performed using the ImageJ software. **F.** Quantification of the mitochondrial area/area occupied by cell as a percentage in control and MS astrocytes. N=10 individual cells per group and quantification was performed using the ImageJ software. **G.** Mitochondrial (*16S*) to nuclear (*RPLP0*) DNA ratio estimative by RT-qPCR. N=3 controls and 3 patients, analyzed in technical triplicates in two independent experiments. Data are represented as the mean ± standard error. Unpaired two-tailed Student’s *t* tests or Mann-Whitney tests were used to compare the groups using GraphPad Prism. * *p* < 0.05; \*\**p* < 0.01.

### MS astrocytes have increased superoxide and proinflammatory chemokine production

As reactive astrocytes can have enhanced reactive oxygen species and cytokine production**^4^**, we assessed superoxide production in mitochondria using Mitosox, and observed a significant increase in the Mitosox MFI in MS astrocytes (Figure 5A), suggesting that these cells have higher superoxide production. We also analyzed cytokine production by the astrocytes and observed increased levels of proinflammatory chemokines in the MS group (Fig 5B to 5E).

**Figure 5.**
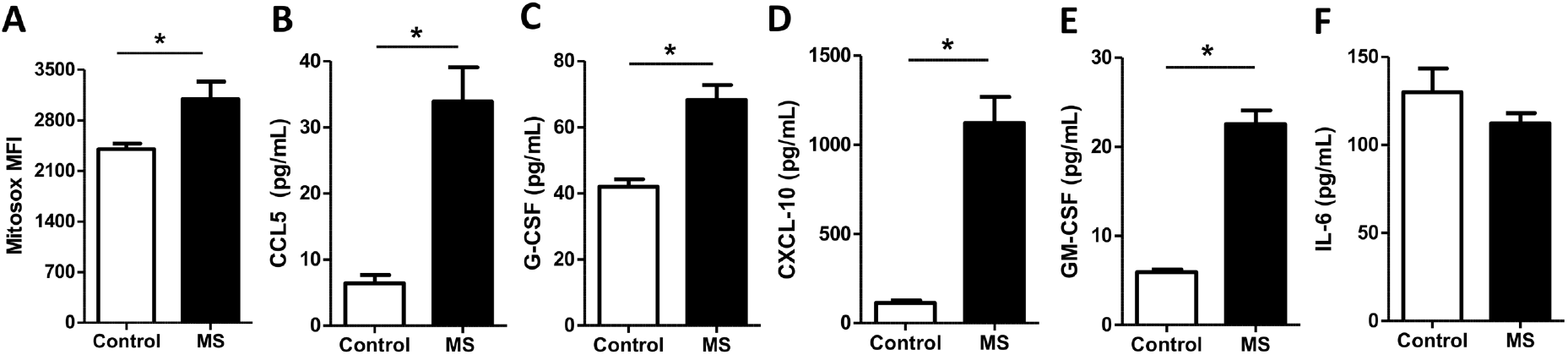
Increased superoxide and chemokine production in MS astrocytes. **A.** Quantification of Mitosox mean fluorescence intensity between the groups. Data are represented as the mean ± standard error. N=3 controls and 3 patients, analyzed in technical duplicates in two independent experiments. Flow cytometry analysis was performed using the FlowJo software. **B-F** Quantification of chemokine and cytokine production using a multiplex assay. Levels of CCL5 **(B)**,G-CSF **(C)**, CXCL-10 **(D)**, GM-CSF **(E)** and IL-6 **(F)** were measured in the supernatant of of 5×10^5^ astrocytes previously activated with 20ng/mL recombinant human TNF-α for 48h. Astrocyte supernatants were collected in two independent experiments. N=3 controls and 3 patients, analyzed in technical duplicates. Data are represented as the mean ± standard error. Unpaired two-tailed Student’s *t* tests or Mann-Whitney tests were used to compare the groups using GraphPad Prism. * *p* < 0.05.

### MS astrocytes display increased electron transport capacity, mitochondrial uncoupling, and proton leak

One of our key objectives in the study was to have deeper insights into the metabolic profile of astrocytes from PwMS. We first performed the Mitostress test (Fig 6A), which quantifies oxygen consumption in intact cells under basal conditions, after the inhibition of mitochondrial ATP synthase by oligomycin (oligo), after maximizing electron transport by the addition of uncoupler (2,4 DNP) and inhibiting the electron transport chain (AA + Rot). We found that non-mitochondrial OCR (Fig 6B), maximal OCR (Fig 6C) and proton leak (Fig 6D) were higher in MS astrocytes, with no differences in ATP-linked OCR (Fig 6E). This indicates enhanced maximal electron transport capacity and mitochondrial uncoupling in these cells, but unchanged basal oxidative phosphorylation-supplied ATP demand. Interestingly, we also observed a significantly increased basal extracellular acidification rate, ECAR (Fig 6F), in MS astrocytes, which could suggest enhanced glycolytic metabolism. We therefore performed the glycolysis stress test (Fig 6G), which quantifies the extracellular acidification after the addition of glucose, after maximizing glycolysis through the inhibition of mitochondrial ATP synthase by oligomycin (oligo) and inhibiting specifically glycolytic metabolism by the addition of 2-DG. We observed that non-glycolytic acidification (Fig 6H) and glycolytic reserve (Fig 6I) rates were higher in MS astrocytes compared to controls, which is suggestive of increased uncoupling, in line with the Mitostress test analysis, and increased cell capability to respond to acutely increased energetic demands, respectively. In fact, astrocytes play several energy demanding processes in MS **^4, 17^**, which highlights the need of increased metabolic plasticity in these cells.

**Figure 6.**
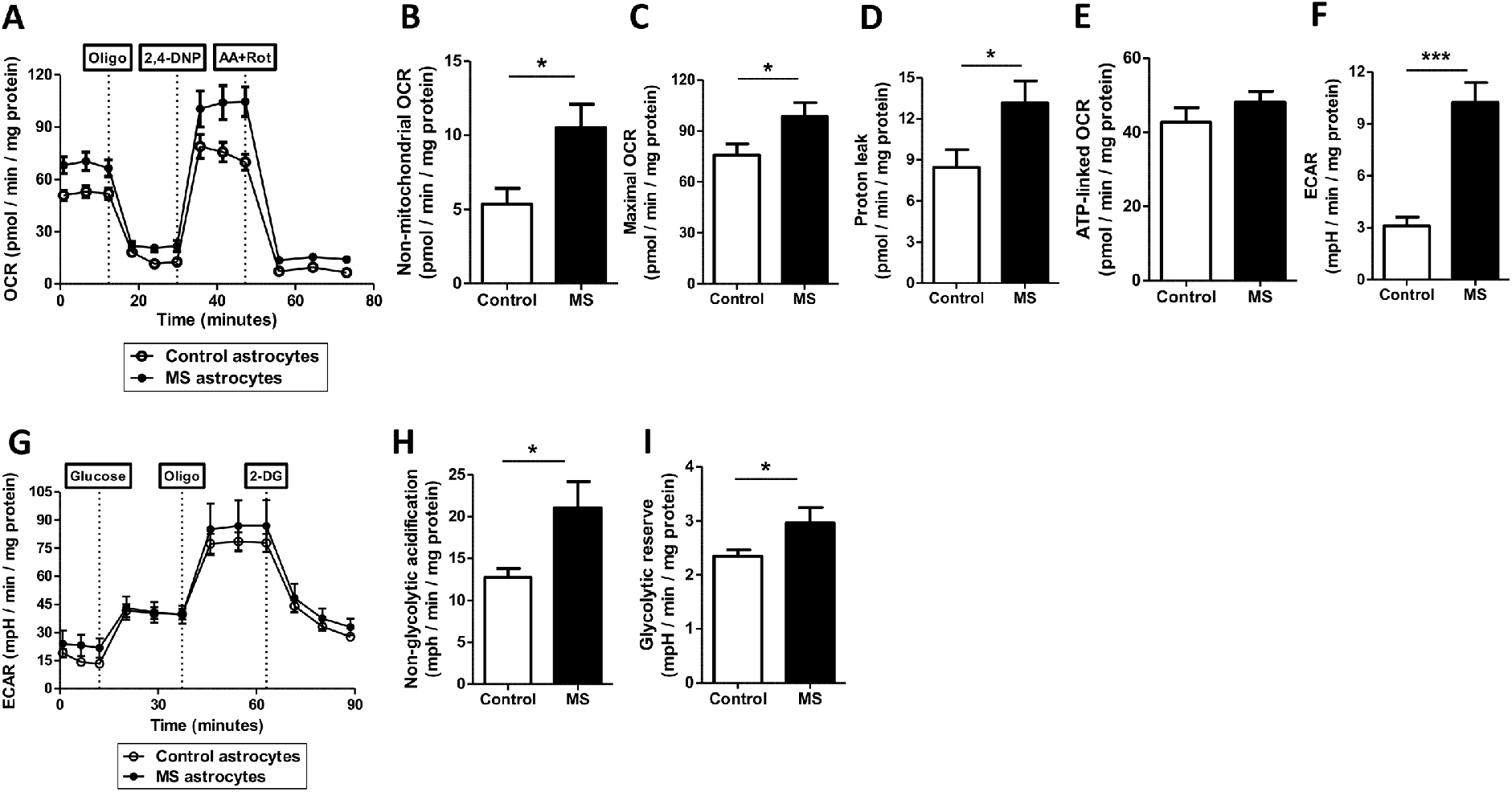
Mitostress and glycolysis stress test assays. **A.** Curves showing the overall OCR for control and MS astrocytes. Oligo: Oligomycin; 2,4-DNP: 2,4-dinitrophenol; AA+Rot – antimycin-A plus Rotenone. **B.** Quantification of non-mitochondrial OCR. **C.** Quantification of maximal OCR. **D.** Quantification of proton leak. **E.** Quantification of ATP-linked OCR. **F.** Quantification of basal ECAR. **G**. Curves showing the overall ECAR for control and MS astrocytes. Oligo: Oligomycin; 2-DG: 2-deoxi-D-glucose. **H.** Quantification of the non-glycolytic acidification. **I.** Quantification of glycolytic reserve rate (glycolytic capacity/glycolysis ratio). Data are representative of N=3 controls and 3 patients, with cells analyzed in triplicates or quadruplicates in two or three independent experiments. Analysis was performed using the Wave software. All data were normalized to the cell protein content. Data are represented as the mean ± standard error. Unpaired two-tailed Student’s t tests or Mann-Whitney tests were used to compare the groups using GraphPad Prism. * *p* < 0.05; \*\*\**p* < 0.001.

### MS astrocytes have metabolic alterations

Finally, to have specific insights on the metabolic pathways being used by the astrocytes in the context of MS, we performed UHPLC-MS metabolomics on these cells. First, using PCA we detected that for both reverse phase (Fig 7A) and HILIC (Fig 7B) modes there were no batch variations in our analysis, as the QC samples grouped well, and the control and MS astrocytes clusters were different from each other. Then, we filtered the metabolites with VIP score > 1.0 and *p*-value < 0.05, obtaining 16 significantly different compounds between the groups for the reverse phase (Fig 7C) and 4 for the HILIC (Fig 7D) mode. To gain biological insights with these data, we performed Metabolite Set Enrichment Analysis (MSEA) on the metabolites \that presented an enriched concentration level (Fig 7E) and on the metabolites with reduced concentration levels (Fig 7F) in MS astrocytes, observing 4 significantly enriched pathways related mainly to amino acid biosynthesis and sphingolipid metabolism, which have already been linked to MS pathology**^18–21^**.

**Figure 7.**
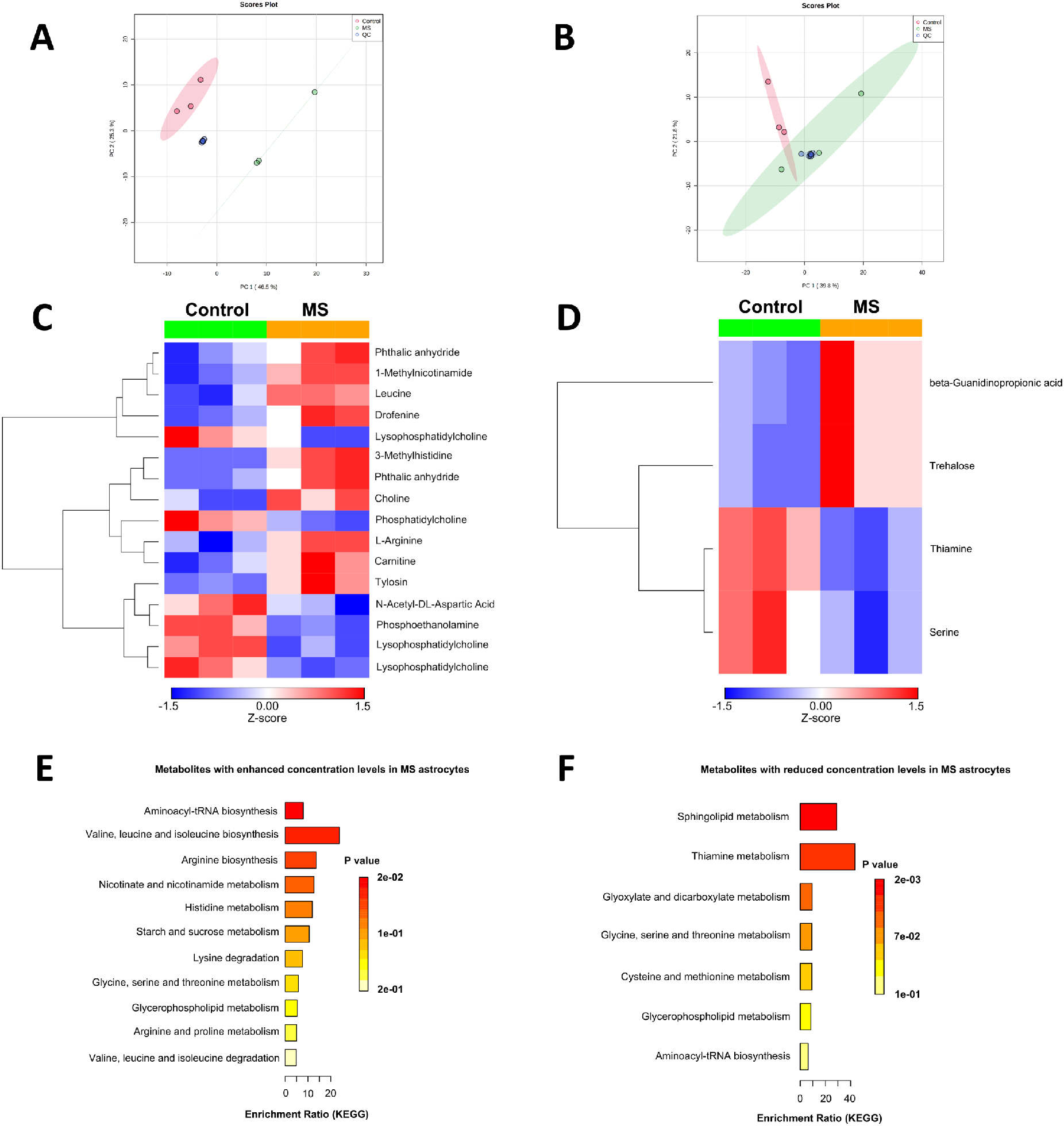
UHPLC-MS metabolomics. **A.** PCA scores plot of the samples in the reverse phase mode. **B.** PCA scores plot of the samples in the HILIC mode. Quality control (QC) samples were included in the PCA analyses to ensure that no batch variations were present in the experiments. **C.** Heatmap representing all 16 metabolites that were significantly different between groups (VIP score > 1.0 and *p*-value < 0.05) in the reverse phase mode. **D.** Heatmap representing all 4 metabolites that were significantly different between groups (VIP score > 1.0 and *p*-value < 0.05) in the HILIC mode. The heatmaps are presented in a row Z-score scale in which red indicates metabolites with enhanced concentration levels and blue indicates metabolites with reduced concentration levels. **E-F** Metabolite Set Enrichment Analysis on the metabolites with enhanced concentration levels in MS astrocytes **(E)** and the metabolites with reduced concentration levels in MS astrocytes **(F)**. The analysis was performed using the KEGG human metabolite database as a source and the colors are scaled by the *p*-value. Data preprocessing, statistical analysis and visualization were performed using MetaboAnalyst software and R programming language. Peak detection, deconvolution, alignment, identification, compound annotation and background subtraction was performed using the MS-Dial software. Compound annotation was performed using LipidBlast and MassBank of North America library for polar metabolites and manually curated to prevent false positives/negatives and only metabolites with a similarity score higher than 70% were selected. After preprocessing, the data were normalized using LOWESS normalization on MS-Dial and the normalized data were exported for statistical analysis. A relative standard deviation cutoff of 30% was applied on the dataset, followed by log2 transformation and autoscaling. *‘Barplot’*, *‘heatmap.2’* and *‘gplot’* libraries and packages were used for data visualization. To observe the differential metabolites between HC and MS samples, variable selection methods were applied. Each dataset was subjected to *t*-test and PLS-DA VIP score analysis using the *‘stats’* and *‘mixOmics’* packages. N=3 controls and 3 PwMS.

## DISCUSSION

MS is a complex disease in which mechanisms are not completely understood. One difficulty in studying the disease is that it is not fully recapitulated by animal models**^9^** and has interspecific differences that make it very hard to translate mouse study findings into humans**^8^**. Here, we used the iPSC technology to derive astrocytes from RRMS patients to find differences that could recapitulate disease aspects, focusing mainly on cellular metabolism and mitochondrial function. We could successfully derive and characterized these astrocytes, in accordance with the few published studies using iPSC-derived cells to model MS**^22–26^**.

To our knowledge, there are only two articles in the literature that analyzed iPSC-derived astrocytes from PwMS, which reached very different conclusions**^25, 26^**. While the first**^25^** suggested that MS alterations should be mostly linked to changes in the immune response rather than those occurring within the CNS, the second**^26^** suggested exactly the opposite, that the resident CNS cells must play a significant role in the development of MS, acting in synergy with peripheral immune cells to promote autoimmune neuroinflammation. Although Ponath et al**^26^** analyzed some metabolic parameters in astrocytes bearing a NF-κB gain of function variant, the authors did not assess cell metabolism in further details, nor mitochondrial functionality and morphology within these cells. It is important to point out that there are some clinical parameters in these studies that may have contributed to the lack of differences between control and MS astrocytes. In the first study**^25^** the samples were all obtained from patients at the time of their first relapses, so the disease progression was at its very early stages. In the second study**^26^**, the authors included RRMS and SPMS patients in the same analysis and all of them were receiving highly effective treatments with monoclonal antibodies, including rituximab and natalizumab. Here we included instead only RRMS patients, some of them with a history of several relapses, that were not under immunosuppressive therapies and whose age was similar to the control group, aiming to avoid possible sources of variation.

Mitochondria can participate in the pathology of several neurologic diseases through modulation of mitochondrial morphology, mitophagy, generation of oxidants, interactions with the endoplasmic reticulum, and regulation of calcium metabolism**^27^**. Importantly, several mitochondrial alterations have already been described in PwMS**^28, 29^**, highlighting the importance of these organelles to the disease pathology. Regarding the genes we found upregulated in MS astrocytes, *UXT* has already been associated with increased NF-κB activity**^30^**. Also, several Solute Carrier 25 (SLC25) transporters have already been suggested as markers for mitochondrial dysfunction in the brain**^31^**. When analyzing the pathways and processes enriched in the MS astrocytes, mitophagy was highlighted along with neurodegeneration and mitochondrial transport, in line with observations in PwMS**^6, 32^**. In this sense, we also observed increased mitochondrial fission in MS astrocytes, which has already been reported to drive neurodegeneration**^16, 33^**.

Accumulation of unhealthy mitochondria in the cells leads to increased superoxide formation and mitochondrial DNA damage**^34^**. Interestingly, we in fact observed in this study significantly increased MitoSOX oxidation, suggestive of higher mitochondrial superoxide levels in MS astrocytes, together with a decreased mitochondrial/nuclear DNA ratio, corroborating these speculations.

When analyzing the production of inflammatory mediators, we observed increased levels of MS-related chemokines**^35–37^**. This supports the hypothesis that MS astrocytes can enhance neuroinflammation in the CNS and act in synergy with immune cells in orchestrating neurodegenerative mechanisms.

Deepening into the metabolic profiling of the cells, our extracellular flux analysis indicated higher non-mitochondrial OCR, maximal OCR and proton leak in MS astrocytes. The first parameter has already been associated with inflammatory enzymes such as the NADPH oxidases and cyclooxygenases**^34^**. The second indicates higher electron transport capacity and the third suggests mitochondrial uncoupling and oxidative stress**^34^**. One possibility to be considered is that mitochondria in neurons and glial cells are constantly exposed to an enhanced flux of ions, such as calcium, which would require the proton gradient and therefore increase oxygen consumption regardless of ATP demand**^34, 38^**. Accordingly, increased Complex IV activity and oxidative stress markers levels were found increased specifically in astrocytes in lesions from PwMS**^39^**. Also in line with our results, it has been demonstrated that the cerebrospinal fluid of PwMS induces increased proton leak in cultured neurons**^40^**. Although our basal ECAR measurements were suggestive of enhanced glycolysis in MS cells at first, when performing specifically the glycolysis stress test we observed an increased non-glycolytic acidification and increased glycolytic reserve rate in MS astrocytes, which indicates that the enhanced acidification rates are not a result of glycolysis itself but are rather due to increased mitochondrial uncoupling and proton leak in these cells.

Finally, our metabolomics analysis indicated a distinct metabolic profile between control and MS astrocytes. Among the different metabolites, we observed an accumulation of amino acids such as leucine and L-arginine in MS astrocytes, together with a decreased concentration of phospholipids and thiamine. Interestingly, when performing MSEA on metabolites accumulated in MS astrocytes, we observed a significant enrichment of two terms: aminoacyl-tRNA biosynthesis and valine, leucine, and isoleucine biosynthesis, which indicates intense metabolic activity in these cells, and a deficiency in amino acid catabolism. Accordingly, amino acid catabolism in MS regulates immune homeostasis in a way that a deficiency in these catabolic reactions, as already reported in patients, is related to an increased release of proinflammatory cytokines and fewer regulatory T-cells**^18^**. Then, when performing MSEA on metabolites that were reduced in MS cells, we observed an enrichment of sphingolipid and thiamine metabolism. Intriguingly, sphingolipid metabolism is a driver of neurodegeneration and has been considered a therapeutic target in neurologic diseases**^19^**. Sphingolipid metabolism has also been shown to induce a proinflammatory phenotype in astrocytes, which drives the pathology in mice**^20, 21^**, although this has never been demonstrated in patients. Specifically in MS, sphingosine-1-phosphate receptor is a therapeutic target of the widely used medication fingolimod, which is approved for the treatment of PwMS, and it has been demonstrated that beyond its canonical role in inhibiting lymphocyte trafficking to the CNS, this drug induced an antiinflammatory phenotype in human astrocytes**^41^**, suggesting that part of the effects of fingolimod may rely on the modulation of astrocyte functions in PwMS, which should be further investigated. Additionally, thiamine (vitamin B1) deficiency was described to induce neurologic manifestations in PwMS, mainly related with depressive behavior**^42^**.

In summary, we described here the metabolic profile of iPSC-derived astrocytes from PwMS, demonstrating that these cells mimic several previously reported disease features and validating this model as a non-invasive tool for future mechanistic and drug targeting studies in MS. A summary of our findings is shown below (Fig 8).

**Figure 8.**
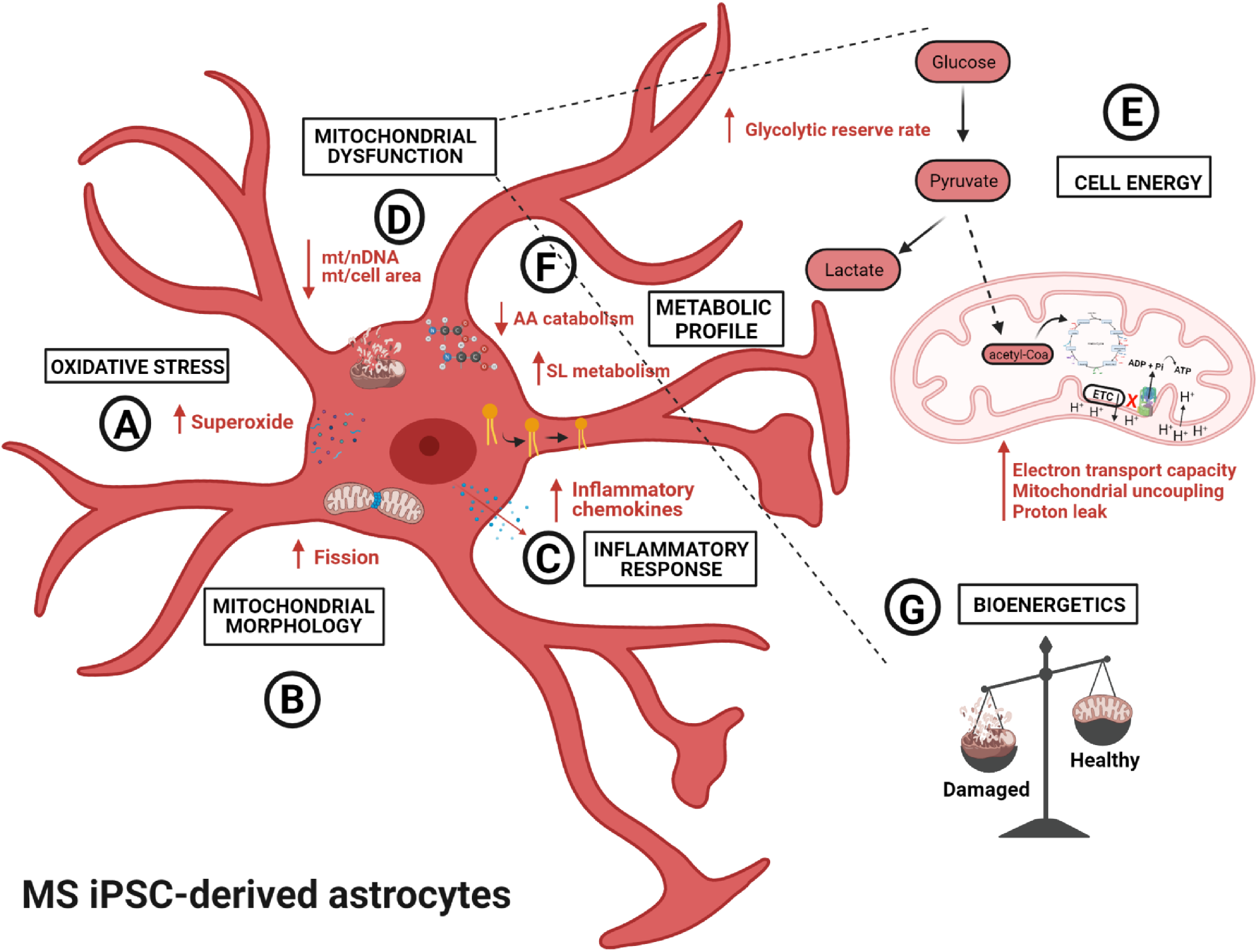
Summary of study findings. In this study, we were able to successfully derive astrocytes from Multiple Sclerosis patients using the iPSC technology. When compared to controls, we observed in these cells changes in oxidative stress, including increased superoxide production **(A)**; changes in mitochondrial morphology, with increased mitochondrial fission **(B)**; changes in the inflammatory response, with an increased secretion of proinflammatory chemokines **(C)**; mitochondrial dysfunction, with decreased mitochondrial/nuclear DNA ratio and mitochondrial/cell area ratio **(D)**. This mitochondrial dysfunction could be demonstrated in metabolic alterations, with enhanced mitochondrial uncoupling, proton leak and electron transport capacity in MS astrocytes, regardless of oxidative phosphorylation-supplied ATP demands. We also observed an increased glycolytic reserve rate in these cells, suggesting increased capability to respond to acutely increased energetic demands **(E)**. We also observed differences in the metabolic profile of the MS astrocytes, with increased sphingolipid metabolism and decreased amino acid catabolism **(F)**. Overall, our results indicate that MS astrocytes have an increased mitochondrial damage, which can enhance the pathology of the disease by impairing the cellular bioenergetic profile and inducing the production of oxidants (**G**). mt/nDNA: mitochondrial to nuclear DNA ratio; mt/cell area: mitochondrial area/area occupied by cell; ETC: electron transport chain; Pi: inorganic phosphate; AA: amino acid; SL: sphingolipid; ATP: adenosine triphosphate; ADP: adenine diphosphate; H^+^: hydrogen protons. The figure was generated using the Biorender software.

## ACKNOWLEDGEMENTS

We thank CEFAP-USP, especially Dr. Iuri C. Valadão and Dr. Mário C. Cruz for their assistance with confocal microscopy analysis. We also thank Dr. Naila C. V. Lourenço from the Human Genome and Stem-Cell Research Center-USP for her assistance with the MLPA analysis. We also acknowledge all the staff from the electron microscopy facility at the School of Medicine-USP.

This work was supported by Fundação de Amparo à Pesquisa do Estado de São Paulo (Grants 2017/05264-7; 2018/23460-0; CEPID: 2013/08028-1; Centro de Pesquisa, Inovação e Difusão em Processos Redox em Biomedicina: 2013/07937-8) and Coordenação de Aperfeiçoamento de Pessoal de Nível Superior (Financial Code 001).

## AUTHOR CONTRIBUTIONS

B.G., D.F.O., A.J.K., E.M.L.O., M.Z. and N.O.S.C contributed to conception and design of the study. B.G., D.F.O., M.C., P.J.B., J.L., C.N.S.B., H.C.R., C.C.C.S., M.I.H., E.G.C., A.S and E.M.L.O. contributed to the acquisition and analysis of data. B.G. and N.O.S.C. contributed to drafting the text and preparing the figures. All authors revised and approved the final version of the manuscript.

## CONFLICTS OF INTEREST

The authors declare no conflicts of interest.

## Notes

### Competing Interest Statement

The authors have declared no competing interest.

